# PepHammer – a lightweight web-based tool for bioactive peptide matching and identification

**DOI:** 10.64898/2026.04.13.718252

**Authors:** Alexander G. B. Grønning, Camilla Schéele

## Abstract

Peptides are gaining increasing attention as therapeutic agents. Already, peptide-based therapeutics play a key role in the treatment of diverse diseases, including diabetes, obesity, and other complex disorders, and their clinical relevance is expected to expand further in the coming years.

Technological and computational advances have substantially enriched peptidomics, massively increasing the scale and depth of peptide identification. As a result, increasingly large and information-rich datasets are now available for downstream analysis and experimental validation. However, the rapid expansion of peptidomics datasets also leads to a corresponding increase in search space, complicating the efficient identification of peptides relevant to specific biological or clinical questions.

To address this challenge, we present *PepHammer*, a lightweight web-based tool for bioactive peptide matching and identification. PepHammer allows users to input up to 10000 peptides (2–150 amino acids in length) and compare them against extensive databases of peptides with predicted or experimentally validated bioactivities and tissue associations using Hamming distance, Grantham distance, as well as partial or exact matching strategies.

Via an example study of human milk peptidomics, we demonstrate that PepHammer rapidly provides an overview of the bioactivity and tissue-relational landscape, serving as a starting point for downstream analyses. PepHammer thus enables efficient exploration of large-scale peptidomics datasets and facilitates the identification of biologically relevant peptides.

## Introduction

Peptides are known signaling molecules, and have emerged as an important class of therapeutic agents due to their high specificity, favorable safety profiles, and broad applicability across disease area. Peptide-based drugs are already widely used in the treatment of metabolic disorders such as diabetes and obesity [1], as well as in areas including oncology, immunology, and neurology. Continued advances in peptide engineering and delivery strategies are expected to further expand their clinical utility, positioning peptides as a central modality in future therapeutic development [2, 3].

In parallel, recent developments in mass spectrometry-based peptidomics and associated computational workflows have dramatically increased the depth and scale of peptide identification [4, 5]. Modern analytical platforms now enable comprehensive and detailed profiling of endogenous peptides, resulting in rapidly growing peptidomics datasets [4–7], which in turn provide unprecedented opportunities to explore peptide bioactivity and function at a large scale.

Despite these advances, the increasing size and complexity of peptidomics datasets present a significant analytical challenge. In particular, the expansion of the candidate search space complicates the prioritization of peptides with potential biological or therapeutic relevance. Existing databases of bioactive peptides and peptide bioactivity prediction tools provide valuable resources [8]; however, efficiently mapping large experimental datasets to known bioactivities and predicting peptide function at scale remain non-trivial and time-consuming tasks, particularly for users without extensive computational expertise. In addition, determining whether peptides of interest have been previously identified in peptidomics datasets from other tissue types remains challenging.

To address this gap, we developed *PepHammer*, a lightweight web-based tool for bioactive peptide matching and identification. PepHammer enables users to screen up to 10000 peptides against bioactivity and human peptidomics tissue databases (**Figure 1**) using similarity metrics such as Hamming distance and Grantham distance [9], and exact or partial sequence matching. By reducing the effective search space and enabling rapid bioactivity and tissue-based annotation, PepHammer facilitates the initial exploration of peptidomics datasets and supports the identification of candidates for downstream functional analysis.

**Figure 1.**
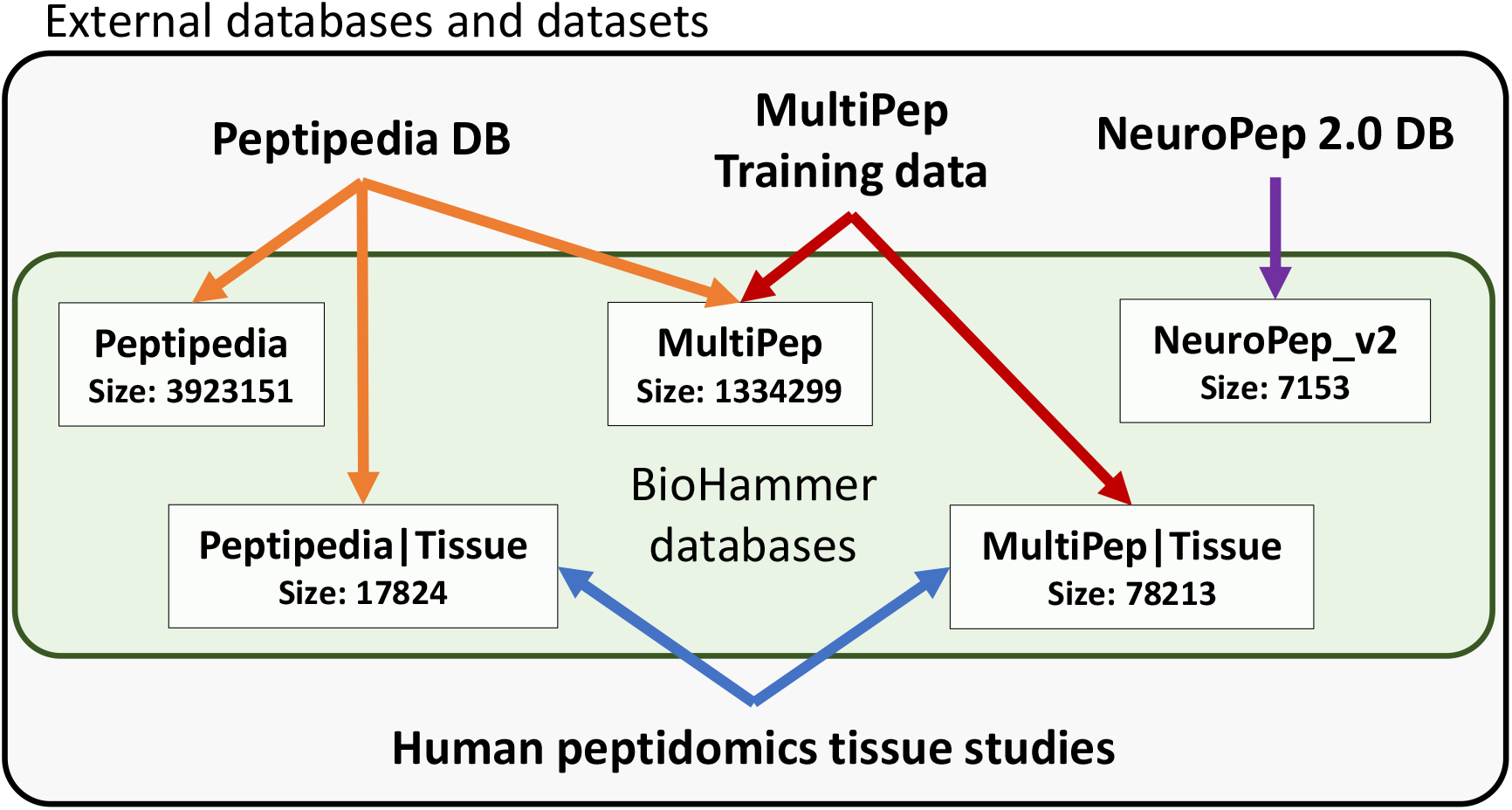
BioHammer databases. Overview of the sizes of the BioHammer databases and their underlying data sources. Human peptidomics tissue studies are from [10–20].

In this work, we present PepHammer, including a detailed overview of its interface, underlying data sources, and implemented methods. The online tool is available here: https://cphbat.shinyapps.io/pephammer/. In addition, we provide an example study of human milk peptidomics, where we demonstrate that PepHammer rapidly can characterizes the bioactivity and tissue-relational landscape, providing a foundation for downstream analyses.

## PepHammer interface

The PepHammer web interface consists of three main tab pages: Pep_Search, Statistics, and Cite. The ‘Pep_Search’ tab enables database queries, the ‘Statistics’ tab provides an overview of database content, and the ‘Cite’ tab contains citation information.

Both the ‘Pep_Search’ and ‘Statistics’ tabs are organized into a left-hand sidebar and a main display area. The sidebar contains adjustable input controls and action buttons, while the main display area presents the corresponding outputs.

### Pep_Search

An example of the ‘Pep_Search’ tab is shown in **Figure 2**. The basic workflow consists of three main steps: (1) input or upload query peptides (**Figure 2**, point 4); (2) select the target database for mapping (**Figure 2**, point 2); and (3) define the mapping method between query and database peptides (**Figure 2**, point 3). The results are presented as an interactive table in the main display area.

**Figure 2.**
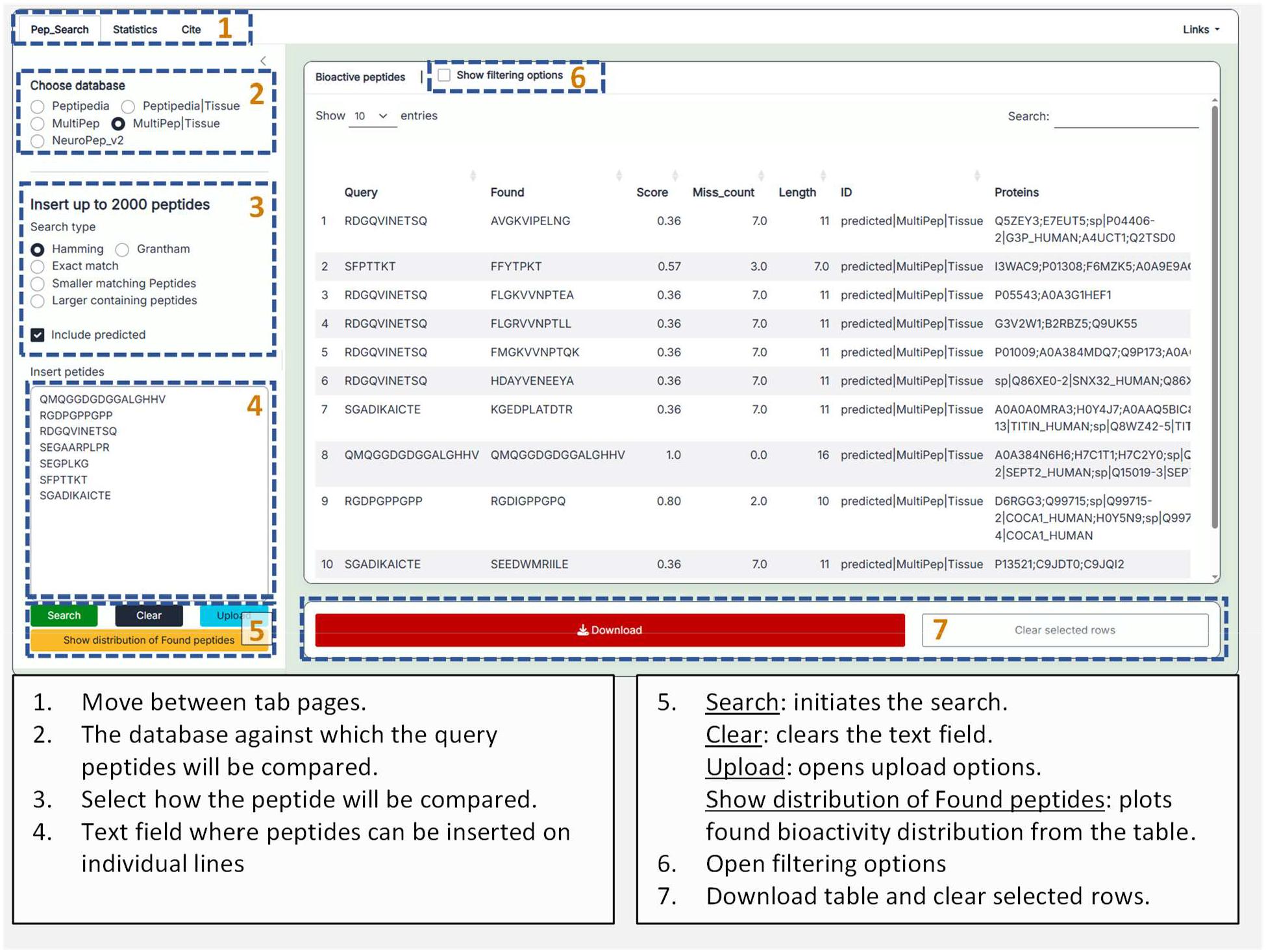
Pep_Search tab page. Dashed dark blue boxes highlight numbered elements corresponding to the descriptions provided in the text boxes below.

#### PepHammer databases

Users can select from the databases listed below (see **Figure 1**). All peptides included have lengths ranging from 2 to 150 amino acids. The following presents as list of the PepHammer databases:

- **Peptidpedia**. This database is based on all bioactive peptides from Peptipedia 2.0 [21]. It contains both peptides with predicted bioactivities and bioactivities integrated from external peptide databases. *ID format:* Peptipedia ID (integer) or Peptipedia ID appended with “|predicted”.
- **Peptipedia**|**Tissue**. This database contains peptides from the PepHammer Peptipedia database that overlap with those identified in the human peptidomics studies. *ID format:* Peptipedia ID (integer) or Peptipedia ID appended with “|predicted”.
- **MultiPep**. This database comprises peptides from the PepHammer Peptipedia database restricted to sequences containing only natural amino acids, with bioactivity predictions generated using the MultiPep tool [22]. Only peptides with at least one predicted bioactivity score > 0.5 are retained. In addition, peptides from the training data of the MultiPep prediction tool are included. *ID format:* “db|MultiPep|” + Peptipedia ID or “predicted|MultiPep|” + Peptipedia ID.
- **MultiPep**|**Tissue**. This database combines peptides from the PepHammer MultiPep database with peptides identified in the human peptidomics studies, all annotated using MultiPep predictions. These peptides are not filtered based on any prediction threshold. *ID format:* “db|MultiPep|Tissue” or “predicted|MultiPep|Tissue”.
- **NeuroPep_v2**. This database contains peptides from NeuroPep 2.0 [23]. No peptide-specific IDs are provided.

For the PepHammer databases ‘Peptipedia’, ‘Peptipedia|Tissue’, and ‘NeuroPep_v2’, bioactivities are annotated in a binary manner (1 indicating membership in a given bioactivity class). In contrast, for the ‘MultiPep’ and ‘MultiPep|Tissue’ databases, bioactivity annotations are provided as continuous scores ranging from 0 to 1.

Detailed information on the human peptidomics studies used as tissue reference can be found in the Methods section (see **Human peptidomics studies**).

All peptides from the PepHammer database have been mapped to reviewed, unreviewed and isoform proteins from the Uniprot database (Date: 18/03/2026) [24].

#### Search type

PepHammer supports the following search types:

- **Hamming:** Compares query peptides to database peptides of identical length and returns those with the minimal Hamming distance.
- **Grantham:** Compares query peptides to database peptides of identical length and returns those with the minimal Grantham distance.
- **Exact match:** Identifies database peptides that are identical to the query peptides.
- **Smaller matching peptides:** Identifies peptides one amino acid shorter than the query peptides that fully match a subsequence of the query.
- **Larger containing peptides:** Identifies peptides one amino acid longer than the query peptides that contain the full query sequence.
- **Include predicted**: If unchecked, only unpredicted peptides from external databases are included.

#### Interactive table and plots

##### Table

The results are presented in an interactive table containing the following columns:

- **Query:** Input query peptides.
- **Found:** Matching peptides identified by PepHammer in the selected database.
- **Length:** Length of the ‘Found’ peptides.
- **ID:** Peptide origin/type (e.g., predicted, derived from external databases, or tissue-associated).
- **Proteins:** Proteins to which the ‘Found’ peptides can be mapped.
- **Bioactivities/Tissue types:** Associated bioactivity classes and tissue types.

Additional columns for Hamming distance searches:

- **Miss_count:** Number of amino acid substitutions required to transform the ‘Found’ peptide into the query peptide (i.e., the Hamming distance).
- **Score:** Similarity score defined as 1 − (Miss_count /Length). A value of 1 indicates an exact match, whereas lower values indicate increasing dissimilarity.

Additional columns for Grantham distance searches:

- **Miss_count:** Number of amino acid substitutions required to transform the ‘Found’ peptide into the query peptide.
- **Distance:** Grantham distance, where lower values indicate greater chemical similarity between peptides.
- **Percent_of_worst:** Ratio of the observed Grantham distance to the maximum possible (i.e., most chemically dissimilar) distance.

##### Plots

The table is interactive, allowing users to sort entries by clicking column headers. In addition, selecting a table row generates a plot displaying the bioactivity classes and tissue types associated with the corresponding *Found* peptide.

When the “Show distribution of found peptides” button (**Figure 2**, point 5) is clicked, a distribution plot summarizing the bioactivity classes and tissue types of all peptides in the table is displayed below the table.

#### Filtering options

An example of the filtering options menu is shown in **Figure 3**. The available sliders and checkboxes enable dynamic filtering of peptides based on user-defined criteria.

**Figure 3.**
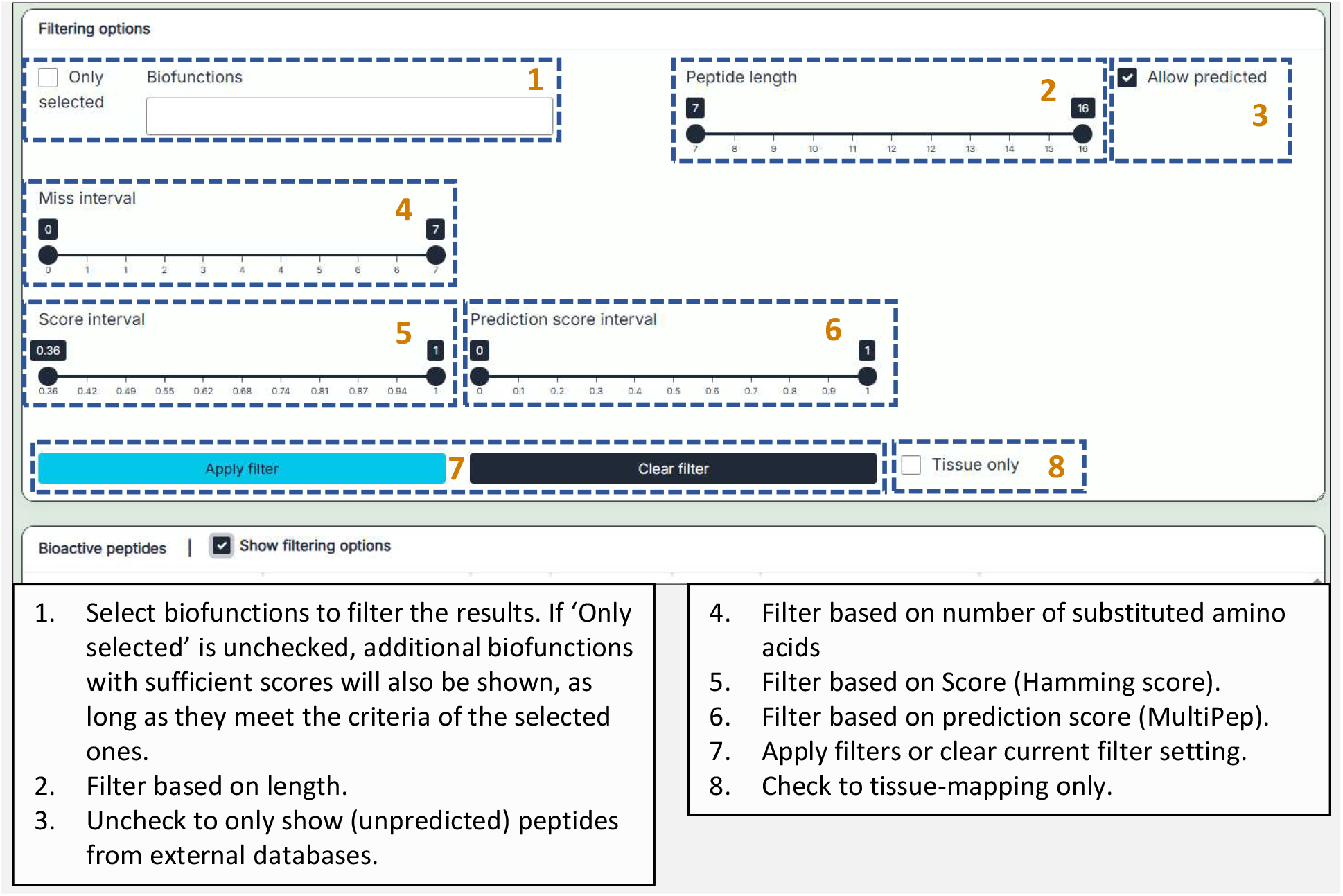
Filtering options. Dashed dark blue boxes highlight numbered elements corresponding to the descriptions provided in the text boxes below.

The filtering menu always includes the following options: ‘Only selected’, ‘Biofunctions’, ‘Peptide length’, and ‘Allow predicted’.

For Hamming distance searches, additional filters are available: ‘Miss interval’ and ‘Score interval’.

For Grantham distance searches, the following filters are provided: ‘Miss interval’, ‘Distance interval’, and ‘Percent_of_worst interval’.

When a tissue-specific database is selected, the ‘Tissue only’ option becomes available, and when any ‘MultiPep’ database has been selected the ‘Prediction score interval’ is available.

### Statistics

The statistics tab provides functions to analyze the PepHammer databases (**Figure 4**). Users can select the desired database and view length, bioactivity and tissue type distributions.

**Figure 4.**
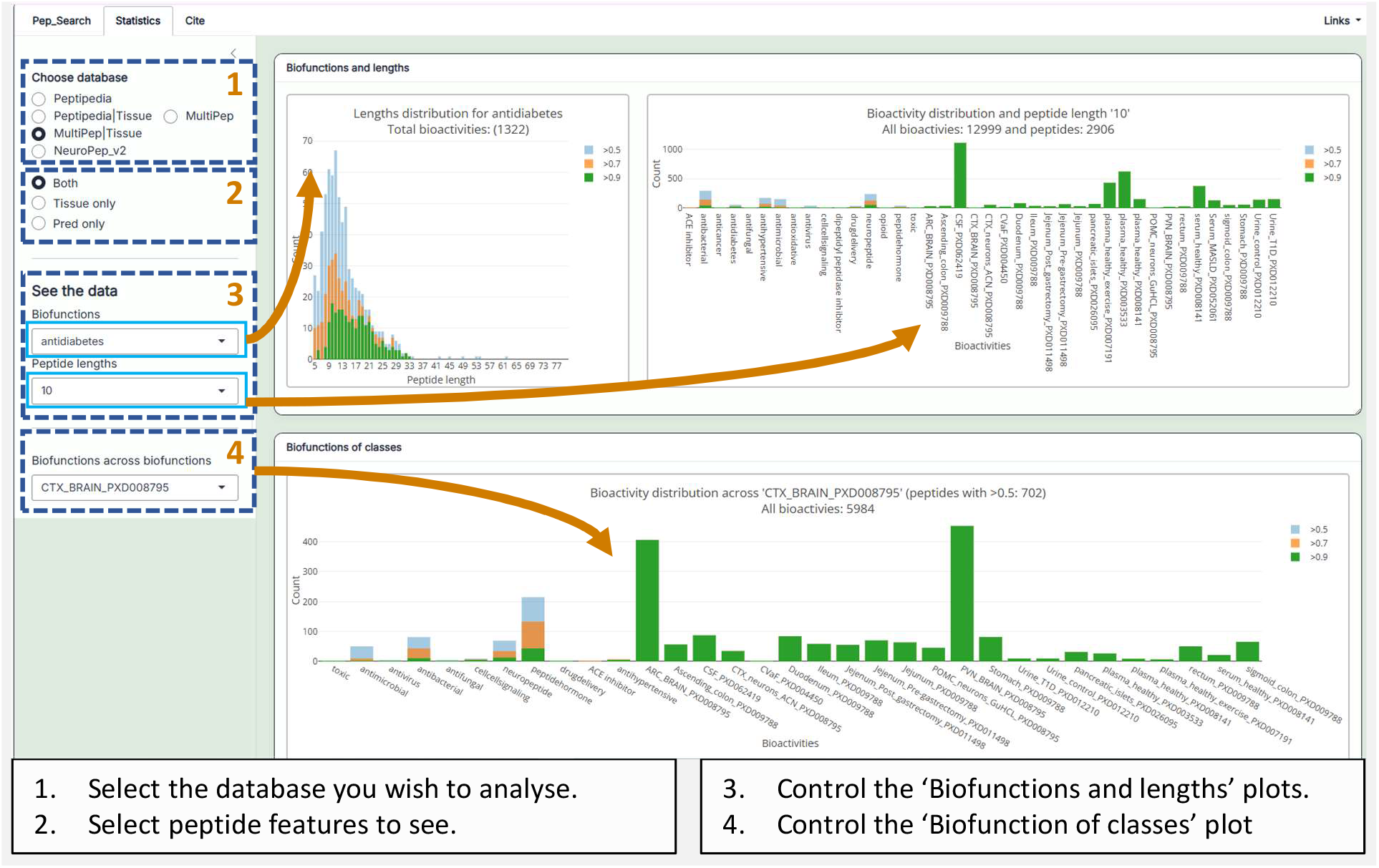
Statistics tab. Dashed dark blue boxes highlight numbered elements corresponding to the descriptions provided in the text boxes below.

## Methods

### Application implementation

The PepHammer webtool is implemented in R v. 4.5.2 [25] using Shiny v. 1.12.1 [26]. Additional applied R packages include: bslib [27], DT [28], PlotLy [29], shinyjs [30], RSQLite [31], DBI [32], future [33], promises [34].

### Hamming distance

To account for ambiguous amino acid codes, Hamming distances were computed using a compatibility matrix defining permissible residue substitutions. A binary matrix was constructed over the set of amino acid symbols, where identical residues were considered fully compatible. In addition, standard ambiguity codes were incorporated such that B was treated as compatible with D and N, Z with E and Q, and J with I and L. The wildcard residue X was considered compatible with all amino acids. During distance calculation, mismatches were only counted when residues were not compatible according to this matrix, thereby extending the conventional Hamming distance to accommodate ambiguous amino acid residue codes.

### Grantham Distance

Grantham distances were computed using the standard amino acid distance matrix, which quantifies physicochemical differences between residues [9]. To accommodate ambiguous amino acid codes, the matrix was extended to include additional symbols. For binary ambiguity codes, distances were defined as the mean of the corresponding canonical residues (e.g., B as the average of D and N, Z of E and Q, and J of I and L). Self-distances for these ambiguous residues were defined as half the distance between their constituent residues. The wildcard residue X was incorporated by assigning distances equal to the mean distance across all canonical amino acids. Pairwise peptide distances were then calculated by summing position-wise Grantham distances across aligned sequences of equal length.

### Human peptidomics studies

Human peptidomics datasets were identified from the PRIDE repository [35]. Peptide sequences were extracted either from mzIdentML (mzID) files using previously published methods [36] or obtained directly from supplementary materials associated with the respective studies. The peptidomics datasets included in this study are listed in **Table 1**.

**Table 1.** PRIDE projects and tissue types. Names are explained if not self-explanatory.

| PRIDE project | Human tissue types |
| --- | --- |
| PXD008795 [10] | ARC_BRAIN: post-mortem mediobasal hypothalamus,<br>CTX_BRAIN: post-mortem cerebral cortex,<br>PVN_BRAIN: post-mortem paraventricular nucleus of the hypothalamus,<br>POMC_neurons_GuHCL: Cultured hypothalamic neurons (HUES9 ES line) treated with guanidine hydrochloride,<br>CTX_neurons_ACN: Cultured cortical neurons (WA09 ES line) treated with acetonitrile. |
| PXD009788 [11] | Ascending_colon, Duodenum, Ileum, Jejunum, Stomach, rectum, sigmoid_colon.<br>All samples were collected during surgery. |
| PXD062419 [13] | CSF: Cerebral spinal fluid from Amyotrophic lateral sclerosis patients and non-neurodegenerative control patients (all peptides were found in both conditions). |
| PXD004450 [14] | CVaF: Cervical-vaginal fluid. |
| PXD011498 [15] | Jejunum_Post_gastrectomy, Jejunum_Pre-gastrectomy |
| PXD052061 [16] | Serum_MASLD: serum from patients suffering from metabolic dysfunction-associated steatotic liver disease. |
| PXD012210 [17] | Urine_T1D: urine from youths with type 1 diabetes,<br>Urine_control: urine from healthy controls. |
| PXD026095 [18] | pancreatic_islets: Pancreatic islets from type 2 diabetes and control patients. |
| PXD003533 [19] | plasma_healthy: plasma from healthy volunteers. |
| PXD008141 [20] | plasma_healthy: plasma from 20 healthy volunteers,<br>serum_healthy: serum from 20 healthy volunteers. |
| PXD007191 [12] | plasma_healthy_exercise: Blood samples from Four healthy male volunteers post-exercise. |

### Example study – bioactivity profile of human milk

To demonstrate the utility of PepHammer as a tool for rapidly generating bioactivity and tissue-association profiles, we analyzed peptides identified in human milk (PXD036477) [37].

Human milk is the sole source of nutrition for many newborns and infants and plays a critical role in health, growth, and development [37]. Although the functions of bioactive peptides in human milk remain incompletely understood, it has been suggested that they contribute to both protective and health-promoting processes [37].

In this example, we generate a bioactivity profile of human milk peptides and provide an overview of the tissues associated with their origin in a non-hypothesis manner. This approach facilitates exploration of human milk peptidomics data, supports hypothesis generation, and enables identification of candidate bioactive peptides, thereby illustrating the applicability of PepHammer.

#### Data extraction

From the PXD036477 repository, we extracted the *peptides*.*txt* file from the *MQresult_PD221-PD576*.*zip* archive. Peptides flagged as “Reverse” or “Potential contaminant” were removed. We further filtered the dataset to retain peptides with non-zero values in at least three LFQ intensity samples, resulting in 8817 unique human milk peptides.

The resulting peptide list was saved as a TXT file (*human_milk*.*txt*), which is available for download from the PepHammer webpage.

#### Results

We applied the MultiPep|Tissue database and searched for exact matches. The resulting bioactivity and tissue profile, based on 988 matching peptides, is shown in Figure 5A. In the unfiltered dataset, bioactivity counts (y-axis) include all predictions with scores > 0.5. Dataset identifiers on the x-axis with the suffix “PXD” correspond to peptidomics tissue datasets.

**Figure 5.**
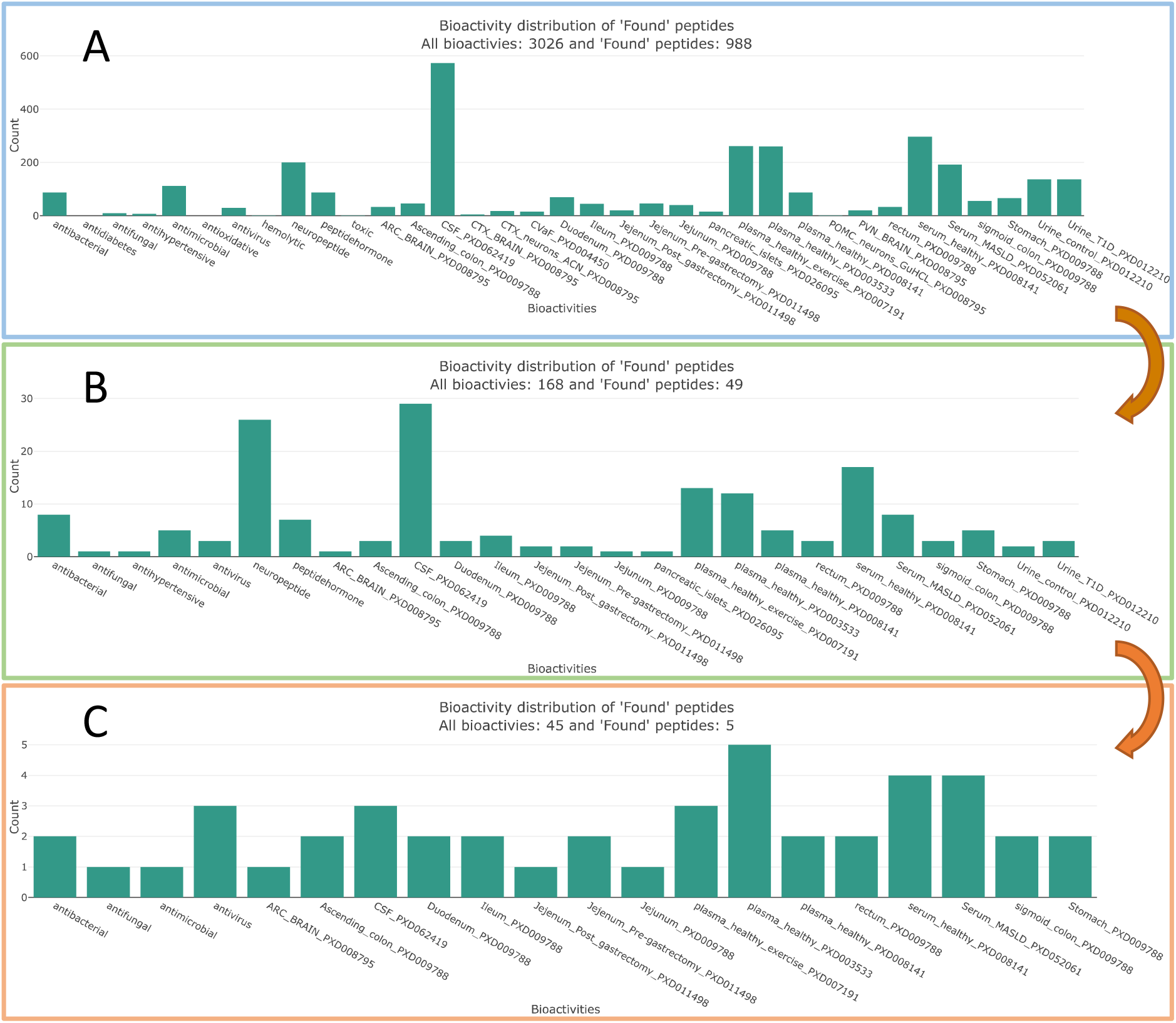
PepHammer bioactivity and tissue profiles. (A) Unfiltered output obtained using the MultiPep|Tissue database. (B) Filtered output after applying a prediction score threshold of ≥ 0.9. (C) Peptides with bioactivities supported by external databases (non-predicted). Orange arrows indicate the filtering steps performed within PepHammer.

A substantial overlap was observed between peptides identified in human milk and those detected in cerebrospinal fluid (573 matches; PXD062419). Additional overlaps were found with plasma datasets (262, PXD007191; 261, PXD003533) and serum datasets (297, PXD008141; 192, PXD052061). Furthermore, 200 peptides were predicted to have neuropeptide activity.

To reduce potential false positives, the prediction threshold was increased to 0.9 (Figure 5B). This filtering step reduced the dataset to 49 peptides while preserving a similar overall profile. Notably, the proportion of predicted neuropeptides increased (26 peptides), and a substantial overlap with the cerebrospinal fluid dataset remained (29 peptides). Corresponding overlaps with plasma and serum datasets were reduced (13, PXD007191; 12, PXD003533; 17, PXD008141; 8, PXD052061).

As a final filtering step, peptides with bioactivity annotations from external databases were identified (Figure 5C). This resulted in five peptides, all of which were mapped to a plasma dataset (PXD003533). These peptides were exclusively annotated as antimicrobial-like.

#### Potential implications of systemic peptide overlap

The observed overlap between human milk peptides and peptides detected in systemic biofluids, including cerebrospinal fluid, plasma, and serum, suggests the presence of a broadly distributed peptide repertoire. As these fluids collectively interface with a wide range of tissues throughout the body, this pattern may indicate that a subset of peptides in human milk reflects conserved or systemically circulating sequences. One possible interpretation is that such peptides could contribute to inter-individual molecular transfer from mother to child. In this context, human milk may serve not only as a nutritional source but also as a vehicle for bioactive peptides with potential roles in early-life physiological development. To this end, it is worth noting the high abundance of predicted neuropeptides, which could play a role in early development. However, this hypothesis remains speculative and requires further experimental validation to determine the origin, stability, and functional relevance of these peptides in the neonatal system.

## Discussion

In this work, we present PepHammer, a lightweight web-based tool for bioactive peptide matching and identification. It enables efficient exploration of large-scale peptidomics datasets and facilitates the identification of biologically relevant peptides by mapping large input sets against databases containing peptides with predicted or experimentally validated bioactivities. In addition, peptides can be mapped to peptidomics datasets across multiple tissue types annotated with bioactivity predictions, enabling cross-tissue comparison and contextualization.

These capabilities support the prioritization of candidate peptides for downstream functional validation and more efficient interrogation of complex datasets. By integrating bioactivity annotation with tissue-specific occurrence and protein identifiers, PepHammer enables systematic linking of peptide sequences to their biological context and functional relevance across tissues.

The databases in PepHammer are derived from multiple external sources, many of which provide additional information about individual peptides. When peptides inherit identifiers from these external databases or datasets, users can consult the original sources to obtain further details. We therefore recommend that users refer to the corresponding external databases whenever relevant peptides are used or analyzed.

In this example study, we analyzed human milk peptidomics data to demonstrate how PepHammer enables rapid annotation of peptides with predicted bioactivities and tissue associations, thereby providing a foundation for downstream analyses and hypothesis generation. In this analysis, exact matching was applied; however, alternative mapping strategies available within PepHammer may offer complementary insights and add further analytical depth.

## Data availabilty

Peptidomics datasets were obtained from the PRIDE database [38] (PXD008795, PXD009788, PXD062419, PXD004450, PXD011498, PXD052061, PXD012210, PXD026095, PXD003533, PXD008141, PXD007191, PXD036477) or from supplementary materials associated with the original publications. PepHammer is available as a web application at: https://cphbat.shinyapps.io/pephammer/.

## Acknowledgements

Novo Nordisk Foundation Center for Basic Metabolic Research is supported by a donation from the Novo Nordisk Foundation (Grant ID number NNF23SA0084103). This project has received funding from the European Research Council under the European Union’s Horizon 2020 research and innovation programme (grant agreement no. 101002725, to C.S.)

## References

1. Clemmensen C, Gerhart-Hines Z, Schwartz TW, et al (2025) Shaping the future of cardiometabolic innovation: advances and opportunities. Nat Metab 2025 78 7:1495– 1497. 10.1038/s42255-025-01343-5

2. Zheng B, Wang X, Guo M, Tzeng CM (2025) Therapeutic Peptides: Recent Advances in Discovery, Synthesis, and Clinical Translation. Int J Mol Sci 2025, Vol 26, Page 5131 26:5131. 10.3390/IJMS26115131

3. Wang L, Wang N, Zhang W, et al (2022) Therapeutic peptides: current applications and future directions. Signal Transduct Target Ther 2022 71 7:48-. 10.1038/s41392-022-00904-4

4. Movassaghi CS, Sun J, Jiang Y, et al (2025) Recent Advances in Mass Spectrometry-Based Bottom-Up Proteomics. Anal Chem 97:4728. 10.1021/ACS.ANALCHEM.4C06750

5. Sachsenberg T, Pino LK, Brunet M, et al (2025) Perspectives in computational mass spectrometry: recent developments and key challenges. Bioinforma Adv 5:vbaf301. 10.1093/BIOADV/VBAF301

6. Yang KL, Yu F, Teo GC, et al (2023) MSBooster: improving peptide identification rates using deep learning-based features. Nat Commun 2023 141 14:4539-. 10.1038/s41467-023-40129-9

7. Demichev V, Messner CB, Vernardis SI, et al (2019) DIA-NN: neural networks and interference correction enable deep proteome coverage in high throughput. Nat Methods 2019 171 17:41–44. 10.1038/s41592-019-0638-x

8. Asim MN, Asif T, Mehmood F, Dengel A (2025) Peptide classification landscape: An in-depth systematic literature review on peptide types, databases, datasets, predictors architectures and performance. Comput Biol Med 188:109821. 10.1016/J.COMPBIOMED.2025.109821

9. Grantham R (1974) Amino Acid Difference Formula to Help Explain Protein Evolution. Science (80-) 185:862–864. 10.1126/SCIENCE.185.4154.862

10. Kirwan P, Kay RG, Brouwers B, et al (2018) Quantitative mass spectrometry for human melanocortin peptides in vitro and in vivo suggests prominent roles for β-MSH and desacetyl α-MSH in energy homeostasis. Mol Metab 17:82–97. 10.1016/J.MOLMET.2018.08.006

11. Dye FS, Larraufie P, Kay R, et al (2019) Characterisation of proguanylin expressing cells in the intestine – evidence for constitutive luminal secretion. Sci Rep 9:15574. 10.1038/S41598-019-52049-0

12. Parker BL, Burchfield JG, Clayton D, et al (2017) Multiplexed Temporal Quantification of the Exercise-regulated Plasma Peptidome. Mol Cell Proteomics 16:2055. 10.1074/MCP.RA117.000020

13. Muqaku B, Dorst J, Wiesenfarth M, et al (2025) Peptidomic analysis of CSF reveals new biomarker candidates for amyotrophic lateral sclerosis. EMBO Mol Med 17:1926. 10.1038/S44321-025-00272-W

14. Muytjens CMJ, Yu Y, Diamandis EP (2017) Discovery of Antimicrobial Peptides in Cervical-Vaginal Fluid from Healthy Nonpregnant Women via an Integrated Proteome and Peptidome Analysis. Proteomics 17:1600461. 10.1002/PMIC.201600461;JOURNAL:JOURNAL:16159861;PAGE:STRING:ARTICLE/CHAPTER

15. Larraufie P, Roberts GP, McGavigan AK, et al (2019) Important Role of the GLP-1 Axis for Glucose Homeostasis after Bariatric Surgery. Cell Rep 26:1399-1408.e6. 10.1016/j.celrep.2019.01.047

16. Mocciaro G, George AL, Allison M, et al (2025) Oxidised Apolipoprotein Peptidome Characterises Metabolic Dysfunction-Associated Steatotic Liver Disease. Liver Int 45:e16200. 10.1111/LIV.16200

17. Van JAD, Clotet-Freixas S, Zhou J, et al (2019) Peptidomic Analysis of Urine from Youths with Early Type 1 Diabetes Reveals Novel Bioactivity of Uromodulin Peptides In Vitro. Mol Cell Proteomics 19:501. 10.1074/MCP.RA119.001858

18. Galvin SG, Kay RG, Foreman R, et al (2021) The Human and Mouse Islet Peptidome: Effects of Obesity and Type 2 Diabetes, and Assessment of Intraislet Production of Glucagon-like Peptide-1. Cite This J Proteome Res 20:4507–4517. 10.1021/acs.jproteome.1c00463

19. Taguchi T, Kodera Y, Oba K, et al (2021) Suprabasin-derived bioactive peptides identified by plasma peptidomics. Sci Rep 11:1047. 10.1038/S41598-020-79353-4

20. Arapidi G, Osetrova M, Ivanova O, et al (2018) Peptidomics dataset: Blood plasma and serum samples of healthy donors fractionated on a set of chromatography sorbents. Data Br 18:1204. 10.1016/J.DIB.2018.04.018

21. Cabas-Mora G, Daza A, Soto-García N, et al (2024) Peptipedia v2.0: a peptide sequence database and user-friendly web platform. A major update. Database 2024:113. 10.1093/DATABASE/BAAE113

22. Grønning AGB, Kacprowski T, Schéele C (2021) MultiPep: a hierarchical deep learning approach for multi-label classification of peptide bioactivities. Biol Methods Protoc 6:bpab021. 10.1093/BIOMETHODS/BPAB021

23. Wang M, Wang L, Xu W, et al (2024) NeuroPep 2.0: An Updated Database Dedicated to Neuropeptide and Its Receptor Annotations. J Mol Biol 436:168416. 10.1016/J.JMB.2023.168416

24. Bateman A, Martin MJ, Orchard S, et al (2023) UniProt: the Universal Protein Knowledgebase in 2023. Nucleic Acids Res 51:D523–D531. 10.1093/NAR/GKAC1052

25. R Core Team (2022) R: A Language and Environment for Statistical Computing

26. Chang W, Cheng J, Allaire JJ, et al (2023) shiny: Web Application Framework for R

27. Sievert C, Cheng J, Aden-Buie G (2026) bslib: Custom “Bootstrap” “Sass” Themes for “shiny” and “rmarkdown”

28. Xie Y, Cheng J, Tan X (2022) DT: A Wrapper of the JavaScript Library “DataTables”

29. Sievert C (2020) Interactive Web-Based Data Visualization with R, plotly, and shiny. Chapman and Hall/CRC

30. Attali D (2021) shinyjs: Easily Improve the User Experience of Your Shiny Apps in Seconds

31. Müller K, Wickham H, James DA, Falcon S (2022) RSQLite: SQLite Interface for R

32. R Special Interest Group on Databases (R-SIG-DB), Wickham H, Müller K (2022) DBI: R Database Interface

33. Bengtsson H (2021) A Unifying Framework for Parallel and Distributed Processing in R using Futures. R J 13:208–227. 10.32614/RJ-2021-048

34. Cheng J (2021) promises: Abstractions for Promise-Based Asynchronous Programming

35. Jones P, Côté RG, Martens L, et al (2005) PRIDE: a public repository of protein and peptide identifications for the proteomics community. Nucleic Acids Res 34:D659. 10.1093/NAR/GKJ138

36. Grønning AGB, Schéele C (2024) Integrating a Multi-label Deep Learning Approach with Protein Information to Compare Bioactive Peptides in Brain and Plasma. Methods Mol Biol 2758:179–195. 10.1007/978-1-0716-3646-6_9

37. Dekker PM, Boeren S, Saccenti E, Hettinga KA (2024) Network analysis of the proteome and peptidome sheds light on human milk as a biological system. Sci Rep 14:7569. 10.1038/S41598-024-58127-2

38. Perez-Riverol Y, Bandla C, Kundu DJ, et al (2025) The PRIDE database at 20 years: 2025 update. Nucleic Acids Res 53:D543–D553. 10.1093/NAR/GKAE1011

